# Rapid diversification of *Pseudomonas aeruginosa* in cystic fibrosis lung-like conditions

**DOI:** 10.1101/231050

**Authors:** Alana Schick, Rees Kassen

**Author notes:** Corresponding author: Alana Schick.

## Abstract

Chronic infection of the cystic fibrosis (CF) airway by the opportunistic pathogen *Pseudomonas aeruginosa* is the leading cause of morbidity and mortality for adult CF patients. Prolonged infections are accompanied by adaptation of *P. aeruginosa* to the unique conditions of the CF lung environment as well as marked diversification of the pathogen into phenotypically and genetically distinct strains that can coexist for years within a patient. Little is known, however, about the causes of this diversification and its impact on patient health. Here, we show experimentally that, consistent with ecological theory of diversification, the nutritional conditions of the CF airway can cause rapid and extensive diversification of *P. aeruginosa.* The increased viscosity associated with the thick mucous layer in the CF airway had little impact on within-population diversification but did promote divergence among populations. Notably, *in vitro* evolution recapitulated patho-adaptive traits thought to be hallmarks of chronic infection, including reduced motility and increased biofilm formation, and the range of phenotypes observed in a collection of clinical isolates. Our results suggest that nutritional complexity and reduced dispersal can drive evolutionary diversification of *P. aeruginosa* independent of other features of the CF lung such as an active immune system or the presence of competing microbial species. They also underscore the need to obtain diverse samples of *P. aeruginosa* when developing treatment plans. We suggest that diversification, by generating extensive phenotypic and genetic variation on which selection can act, may be a key first step in the transition from transient to chronic infection.

**Significance Statement:** Chronic infection with the opportunistic pathogen *Pseudomonas aeruginosa* is the leading cause of lung transplant or death in cystic fibrosis patients. *P. aeruginosa* diversifies in the CF lung, although why this happens remains a mystery. We allowed *P. aeruginosa* to evolve in the laboratory under a range of conditions approximating the CF lung. The diversity of evolved populations was highest, and most closely resembled the range of phenotypes among clinical isolates, in environments resembling the spectrum of nutritional resources available in the CF lung. Our results point to the nutritional complexity of the CF lung as a major driver of diversification and they suggest that diversity could be important in the development of chronic infections.

## Introduction

*Pseudomonas aeruginosa* is a globally ubiquitous, metabolically flexible Gram-negative opportunistic pathogen. While it is an important nosocomial pathogen causing a range of acute infections, *P. aeruginosa* is also commonly recovered from the airways of adult cystic fibrosis (CF) patients where it causes chronic endobronchial infections in up to 80% of adult patients and is the leading cause of morbidity and mortality in this population (Rajan and Saiman 2002, Schaedel et al 2002, Hoiby and Pressler 2006). The majority of chronic CF lung infections are thought to be the result of colonization by *P. aeruginosa* from environmental sources, although highly transmissible epidemic strains are responsible for ~25% of infections in Canadian patients (Aaron et al 2010). Once established, chronic infections can remain persistently associated with a host for decades, being virtually impossible to eradicate with standard antibiotic therapy (Gibson et al 2003).

The transition from environmental strain to one that causes chronic infection is characterized by a few repeatable phenotypic changes underlain by a much larger suite of genetic changes. Free-living environmental strains are typically motile, virulent, and nonmucoid whereas strains isolated from chronically infected CF patients are often non-motile, avirulent, mucoid, and highly antibiotic resistant (Poole 2005, Smith et al 2006, Ciofu et al 2010, Mowat et al 2011, and Sousa and Pereira 2014). These phenotypes represent parallel adaptations to a range of CF lung-specific stressors including increased viscosity, osmotic stress, low oxygen, high concentrations of antibiotics and immune system attack. This parallelism notwithstanding, the other striking feature common to *P. aeruginosa* isolates from CF patients is their diversity. Isolates from within (Jorth et al 2015, Darch et al 2015, Fothergill et al 2010, Mowat et al 2011, Ashish et al 2013, Workentine et al 2013, Clark et al 2015) and among (Dettman et al 2013, Williams et al 2015) patients can be highly diverse, both phenotypically and genetically, making it difficult to identify reliable markers for the onset of chronic infection (Winstanley et al 2016) and predict clinical outcomes on the basis of single colony isolates alone. Indeed, the lack of correlation between the prevalence of a particular phenotype and clinical outcomes (Workentine et al 2013) hints that the diversity of the population, rather than the abundance of any particular phenotype, could be what makes CF lung infections by *P. aeruginosa* so recalcitrant to treatment (Darch et al 2015).

Diversity in the CF lung typically evolves rapidly and *de novo* following colonization (Folkesson et al 2012, Workentine et al 2013, Jorth et al 2015, Darch et al 2015), with genetically and phenotypically diverse clones often persisting for years within the same host (Ciofu et al 2012, Feliziani et al 2014, Sousa and Pereira 2014). This dynamic has many of the hallmarks of an adaptive radiation, the rapid diversification of a lineage into a range of niche specialist types. Both theory (Lewontin 1974, Whittaker and Levin 1975, Nevo 1978) and experiment (reviewed in Futuyma and Moreno 1988, Schluter 2000, Kassen 2009, Kassen 2014) suggest that adaptive diversification is most likely to occur when divergent selection, imposed by ecological opportunities in the form of vacant niche space or underutilized resources, is strong relative to the rate of dispersal. The extent of phenotypic divergence may be further exaggerated, or its rate accelerated, through ecological interactions such as resource competition or predation (Felsenstein 1981, Rozen and Lenski 2000, Doebeli and Dieckmann 2000, Friesen et al 2004).

We suspect that the complex ecological conditions of the CF lung are likely to promote diversification. The CF airway contains a rich spectrum of resources and nutrients that could provide ample ecological opportunity to drive specialization (Palmer et al 2007, Workentine et al 2013, Marvig et al 2014) and the thick mucus layer, a by-product of the impaired ability to transport chloride ions resulting from mutations in the cystic fibrosis transmembrane conductance regulator (CFTR) gene, serves to reduce dispersal among subpopulations. Dispersal will be further reduced due to the highly compartmentalized nature of the lung itself, with distinct patches or habitats such as right and left lobes, upper and lower respiratory tract, and the alveoli and bronchioles on different branches of the respiratory tree. The resource profile and spatial structure of the CF lung thus provide ideal conditions for strong divergent selection to drive diversification. Additional aspects of life in the CF lung such as the emergence of hypermutator lineages (Oliver et al 2000, Feliziani et al 2014), high levels of antibiotic use (Wright et al. 2013), and interactions between *P. aeruginosa* and co-infecting species (Sibley et al 2008, Korgoankar et al 2013), bacteriophages (Brockhurst et al 2005, James et al 2015), and the host immune system (Jensen et al 2010, McKeon et al 2010) may further contribute to the genetic and phenotypic variability seen among isolates from chronically infected patients.

Here, we evaluate the contributions of resource complexity and reduced dispersal to *P. aeruginosa* diversification by tracking the extent of phenotypic diversification within and among independently evolved populations descended from *P. aeruginosa* strain Pa14 after ~220 generations in environments that varied in how closely they resemble the resource profile and viscosity of the CF lung. The most CF-like environment consisted of the nutritionally complex synthetic cystic fibrosis medium (SCFM), a defined medium based on the free amino acid, anion, cation, and carbon source profiles from CF sputum (Palmer et al 2007), supplemented with mucin, the major protein component of mucus, to mimic the thick mucus layer in the lumen of CF patients. Several studies have shown that the presence of mucin increases the viscosity of the media and reduces the motility of *P. aeruginosa* (Wang et al 1996, Landry et al 2006, Fung et al 2010) and is therefore very likely to reduce dispersal. The least CF-like environment was composed of a minimal medium supplemented with glucose as the sole carbon source in the absence of mucin. The experiment comprised factorial combinations of two levels of nutrient complexity (minimal medium versus SCFM) and viscosity (with or without mucin). Phenotypic divergence among independently evolved replicate populations provided an estimate of the contribution of spatial compartmentalization to diversification. We scored both colony morphology and ten traits thought to be associated with patho-adaptation tied to chronic infection of the CF lung (growth rate in LB, pyocyanin and pyoverdine production, biofilm formation, swim and twitch motility, and resistance to four classes of antibiotics: ciprofloxacin, ceftazidime, colistin, and tobramycin) for 12 randomly sampled isolates from each of 12 independently evolved populations within each treatment (for a total of 576 isolates) as well as a collection of 24 *P. aeruginosa* isolates from the lungs of CF patients from across Ontario (Dettman et al 2013) to gauge the extent to which our experimental conditions recapitulate the markers of phenotypic divergence associated with clinical strains.

## Results and Discussion

### Markers of chronic infection evolve in the most lung-like environments

Patho-adaptation to the CF lung is often accompanied by a suite of phenotypic changes that include loss of virulence factors, loss of motility, and increased biofilm formation. We determined the extent to which similar phenotypic changes evolved in our *in vitro* experiment by scoring a range of traits associated with patho-adaptation and using a linear mixed effects model with media and mucin as fixed effects and population as a random effect to examine the impact of different treatment combinations on trait evolution. Our results are summarized for all traits in Table 1 (and SI Appendix, Table S1) and for four traits commonly associated with pathoadaptation in Figure 1.

**Table 1.** Summary of phenotypic evolution. Arrows represent significant deviations from the ancestral value in each treatment group, Bonferroni corrected for multiple comparisons. See SI Appendix Table S1 for test statistics and P-values.

| Trait | SCFM | MIN | SCFM +<br>mucin | MIN +<br>mucin |
| --- | --- | --- | --- | --- |
| Growth rate | — | — | — | — |
| Pyoverdine | ↓ | ↓ | ↓ | ↓ |
| Pyocyanin | — | — | ↑ | — |
| Biofilm | ↓ | ↓ | ↑ | ↓ |
| Swim motility | ↓ | ↓ | ↓ | ↓ |
| Twitch motility | ↓ | ↓ | ↓ | ↓ |
| MIC ciprofloxacin | ↑ | — | ↑ | ↑ |
| MIC ceftazidime | — | — | ↑ | ↑ |
| MIC colistin | — | — | — | ↓ |
| MIC tobramycin | — | — | — | — |

**Figure 1.**
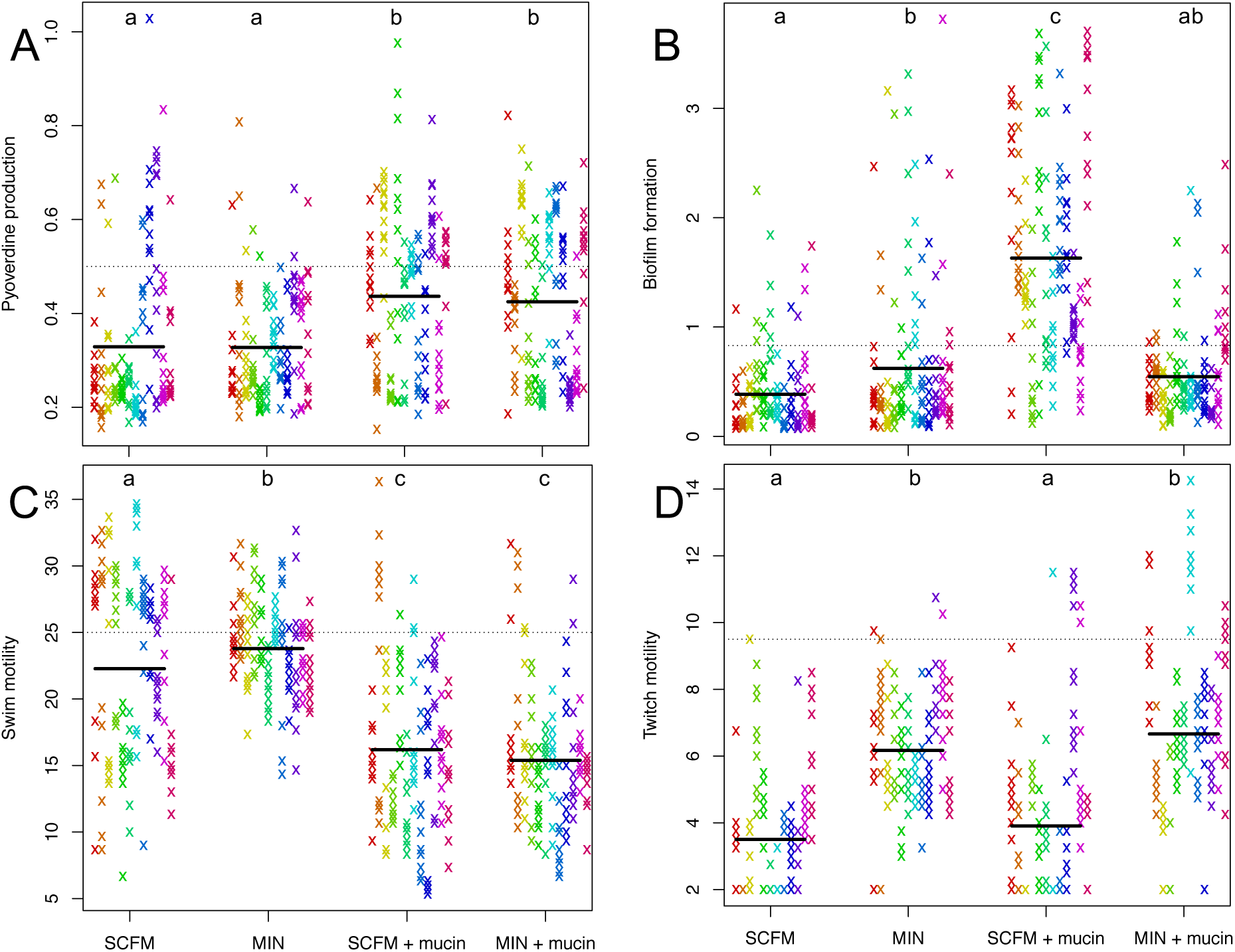
Phenotypic adaptation of evolved isolates for pyoverdine production (A), biofilm formation (B), swim motility (C) and twitch motility (D). Within each treatment, each column/colour represents a replicate evolved population. Solid lines represent treatment means and dashed lines represent the value of that phenotype in the ancestral strain (Pa14).

We observed marked trait evolution including a reduction in pyoverdine production (the main siderophore and a measure of iron-scavenging ability; Fig. 1A), swim motility (Fig. 1C), and twitch motility (Fig. 1D), and an increase in biofilm production (Fig. 1B) in the most CF-like conditions, all changes that mirror those seen during patho-adaptation. These phenotypic changes persist when colonies are subcultured, suggesting they have a genetic basis. Given that our experiment was conducted *in vitro* without competing microflora or an active immune system, these results suggest that selection driven by the combined effects of resource complexity and viscosity is sufficient to explain changes in these patho-adaptive traits characteristic of chronic *P. aeruginosa* infections of the CF lung.

This interpretation might be questioned on the grounds that many of these trait changes, especially those involving a decrease in trait value relative to the common ancestor (pyoverdine production, swim motility, and twitch motility), reflect general adaptation to *in vitro* culture conditions rather than specific adaptation to CF-like environments. This explanation cannot, however, explain the treatment-specific changes in these and other traits (SI Appendix, Table S1). Pyoverdine production, for example, declines less in the presence of mucin than in its absence, while swim motility shows the reverse trend. Changes in twitch motility, on the other hand, were independent of mucin and decreased more in the nutrient rich conditions of SCFM than in minimal media. Pyocyanin and biofilm production both increased in the most CF lung-like conditions (SCFM+mucin) with either no change (pyocyanin) or decreases (biofilm) in the other treatments. Interestingly, the increase in pyocyanin, a virulence factor, is surprising and suggests that the commonly observed decrease in pyocyanin in isolates from CF infections evolves as a response to an aspect of ecology we did not investigate, immune system attack or competition from other species being the most obvious candidates. Taken together, such treatment-specific responses to selection suggest that our results are not simply a by-product of prolonged growth in laboratory culture but, rather, represent trait changes specifically associated with CF lung-like conditions.

A second feature of our results deserves mention here. A lack of evolutionary change in resistance to antibiotics such as colistin and tobramycin is unsurprising, as no drugs were used in the selection experiment. More notable, then, is the marked increase in ciprofloxacin resistance across all conditions except the simplest environment (MIN) and an increase in ceftazidime resistance in environments containing mucin (Table 1). While resistance is commonly observed following antibiotic treatment in clinical settings, our results suggest that both ciprofloxacin and ceftazidime resistance can evolve as a pleiotropic by-product of adaptation to CF lung-like conditions. Spontaneous resistance to antibiotics has been shown to arise via mutations to the MexGHI-OpmD efflux pump that are also associated with biofilm production (Sakhtah et al 2016). This mechanism seems unlikely to explain the increases in resistance in our experiment because we did not observe the expected positive relationship between biofilm formation and resistance for either drug class, the correlation being significantly negative for ciprofloxacin and indistinguishable from zero for ceftazidime (SI appendix, Fig S5). Interestingly, a recent study found that resistance to ceftazidime arose in the absence of this antibiotic as a result of an increase in beta-lactamase production (Davies et al 2017), a mechanism that warrants further exploration.

We also did not observe any mucoid individuals in our evolved populations, which is surprising because mucoidy is a common phenotype of lung-evolved isolates and can often be associated with aggregation and biofilm formation. It is not yet clear why this phenotype did not evolve. If mucoidy represents an adaptation to prolonged infection it is possible that we would have observed it in a longer experiment. An alternative explanation is that mucoidy evolves due to selection for some other factor not captured in our experiment. The mucoid colony morphology is often linked to the overproduction of alginate (Marvig et al 2015), a compound that can also effectively scavenge reactive oxygen species (Simpson et al 1989, Hassett et al 2009). If mucoidy is a pleiotropic result of adaptation for tolerance to oxidative burst from macrophages (Dettman et al 2013) we would not expect to see it evolving in our experiment.

### Within-population phenotypic diversity is driven by resource complexity

We observed extensive phenotypic diversity in all evolved populations. To capture the general trends associated with multivariate phenotypic diversification within and among populations, we used all ten traits described above to calculate the Euclidean distance between pairs of isolates within a population (Eq. 1) and between the means of pairs of populations within each treatment (Eq. 2) as follows:

For isolates, 
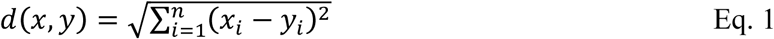

For populations, 
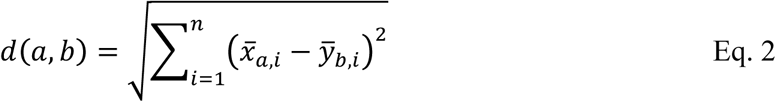

Here, *x_i_* and *y_i_* are standardized *z*-scores for trait *i* of isolates *x* and *y* respectively, and *x_a,i_* and *y_b,i_* are standardized *z*-scores for trait *i* of isolates in populations *a* and *b* respectively. Note that calculating among-population distance using the population means controls for any effect of variation within populations on the measure of among-population distance.

Our results, shown in figure 2, are striking. Populations evolved in the complex nutritional environment of SCFM supported higher amounts of within-population diversity than those evolved in the simpler nutritional environment of minimal media with glucose as the sole carbon and energy source (Fig. 2A; two-factor ANOVA, F_1,44_ = 10.21, p = 0.003), independent of the presence of mucin (F_1,44_ = 0.65, p = 0.423). Moreover, we found no significant interaction between resource complexity and mucin (F_1,44_ = 0.79, p = 0.786). Our results contrast with the prevailing view that the within-host diversification by *P. aeruginosa* accompanying chronic infection is driven primarily by restricted dispersal due to the thick mucus layer in the lung lumen. In evolutionary terms, the nutritional complexity of the lung likely represents abundant ecological opportunity for colonizing *P. aeruginosa* that generates strong divergent selection leading to phenotypic divergence. Competition for resources among incipient niche specialists in the evolving population could further accelerate phenotypic diversification and contribute to coexistence, perhaps through negative frequency-dependent selection (reviewed in Rainey et al 2004, Kassen 2014), leading to long-term persistence of phenotypically divergent *P. aeruginosa* variants within a single host.

**Figure 2.**
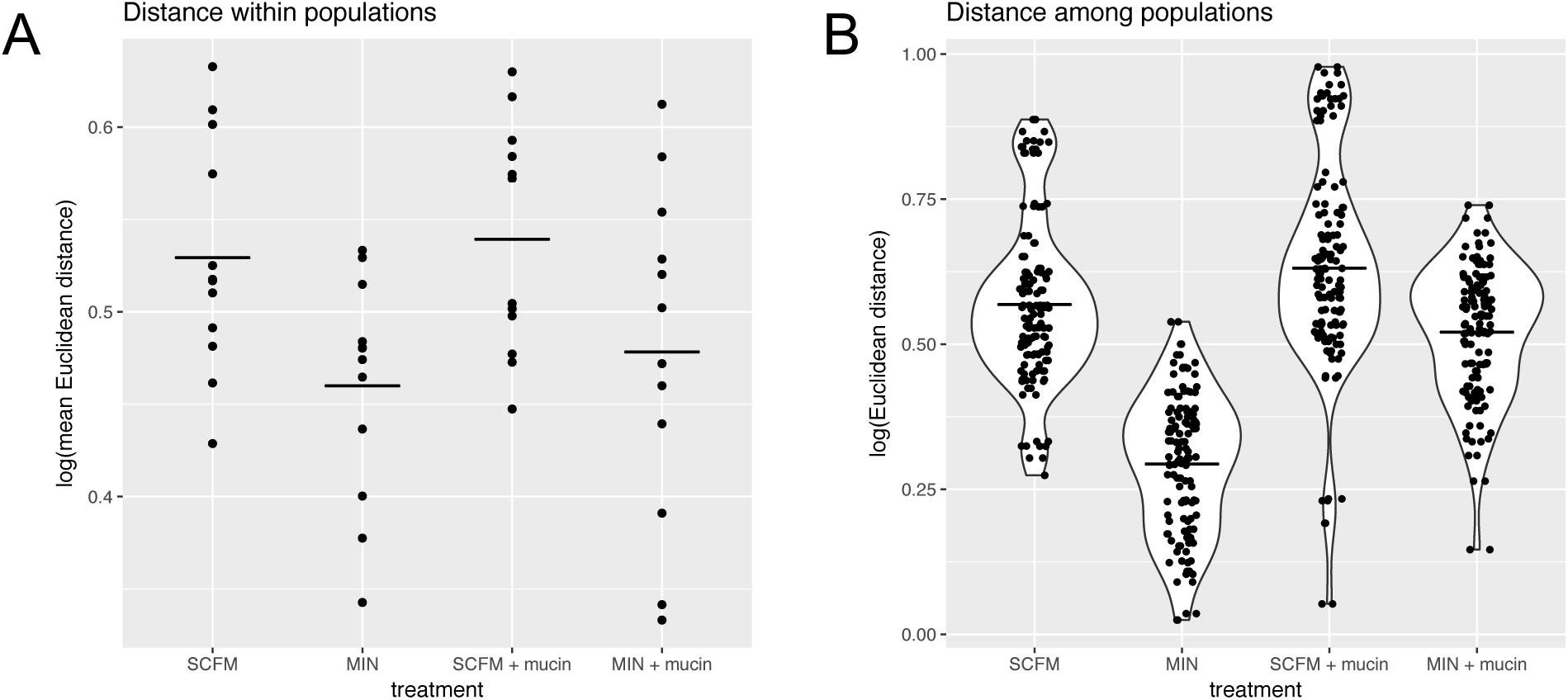
Euclidean distance within (A) and among (B) populations. A) Each point represents a population with the distance within that population determined by the average of all pairwise comparisons of individuals within that population. B) Each point represents the distance between the mean values of two populations. Solid lines represent treatment means in both panels. All distance calculations are Euclidean distance based on all 10 *z*-score transformed phenotypic traits.

### Nutritional complexity and spatial structure cause populations to diverge

*P. aeruginosa* isolates from distinct, spatially separated compartments within the CF airway are often genetically and phenotypically distinct, a result that has been taken to imply niche-specific adaptive diversification following colonization (Markussen et al 2014, Jorth et al 2015). An alternative interpretation is that spatial separation of subpopulations in distinct lung compartments leads to genetic and phenotypic divergence independent of niche-specific adaptation due to stochastic effects in the timing and nature of mutations contributing to pathoadaptation. Our results allow us to quantify the extent of phenotypic divergence among replicate evolved populations within each treatment in our experiment as a proxy for the extent of among-compartment divergence. Both nutritional complexity and mucin had statistically significant effects on among-population divergence (randomization test where treatment labels are resampled and a null distribution of F-statistics calculated; P-values were < 0.0001, <0.0001, and 0.0048 for media, mucin and the interaction, respectively, from a two-factor ANOVA), with the SCFM + mucin treatment displaying the most extensive between-population diversity, followed by the SCFM and MIN + mucin treatments, and the MIN treatment (Fig. 2B). Since replicate populations within a treatment evolved, by design, under the same selective pressure, these results lend support to the idea that mutation-order effects in the most CF-like conditions can support substantial among-population divergence of multivariate phenotypes.

### Comparison of evolved populations and clinical isolates from CF patients across Ontario

To what extent do the conditions in our experiment capture the range of variation associated with *P. aeruginosa* isolates from CF patients? To answer this question, we measured the same ten phenotypic traits of 24 strains of *P. aeruginosa* isolated from the lungs of CF patients from across the Canadian province of Ontario and includes both non-epidemic and epidemic strains (Aaron et al 2010). We then calculated the Mahalanobis distance between each clinical strain and the multivariate distribution of all evolved isolates from each treatment group, separately, to obtain a measure of phenotypic similarity between the clinical strains and the isolates evolved under different conditions. MIN evolved isolates were the most different (largest Mahalanobis distance) from the clinical strains (Fig. 3, ANOVA, F_3, 92_ = 17.9, *P* < 0.001). The clinical strains were most similar to those evolved in SCFM, but not statistically more similar than those evolved in SCFM + mucin or MIN + mucin, evaluated by post-hoc pairwise t-tests, Holm-adjusted for multiple comparisons. Similar results can be seen using a principal component analysis (SI Appendix, Fig. S3). Interestingly, strains classified as epidemic were consistently less similar to lab-evolved isolates than those classified as non-epidemic (two-factor ANOVA, effect of epidemic/non-epidemic, F_1,88_ = 6.21, P = 0.015), a difference largely attributable to levels of antibiotic resistance. Excluding antibiotic resistance traits from the analysis eliminates the difference in Mahalanobis distance between epidemic and non-epidemic strains from our *in vitro* isolates (SI Appendix, Fig. S2, two-factor ANOVA, effect of epidemic/non-epidemic, F_1,88_ = 1.69, p = 0.197). This result further supports those noted by Dettman et al (2013) that the key distinguishing feature of epidemic strains is that they are more resistant to antibiotics than nonepidemic strains. Importantly, epidemic and non-epidemic strains are indistinguishable in the other phenotypic dimensions measured here making it difficult to distinguish them on the basis of non-antibiotic phenotypes alone.

**Figure 3.**
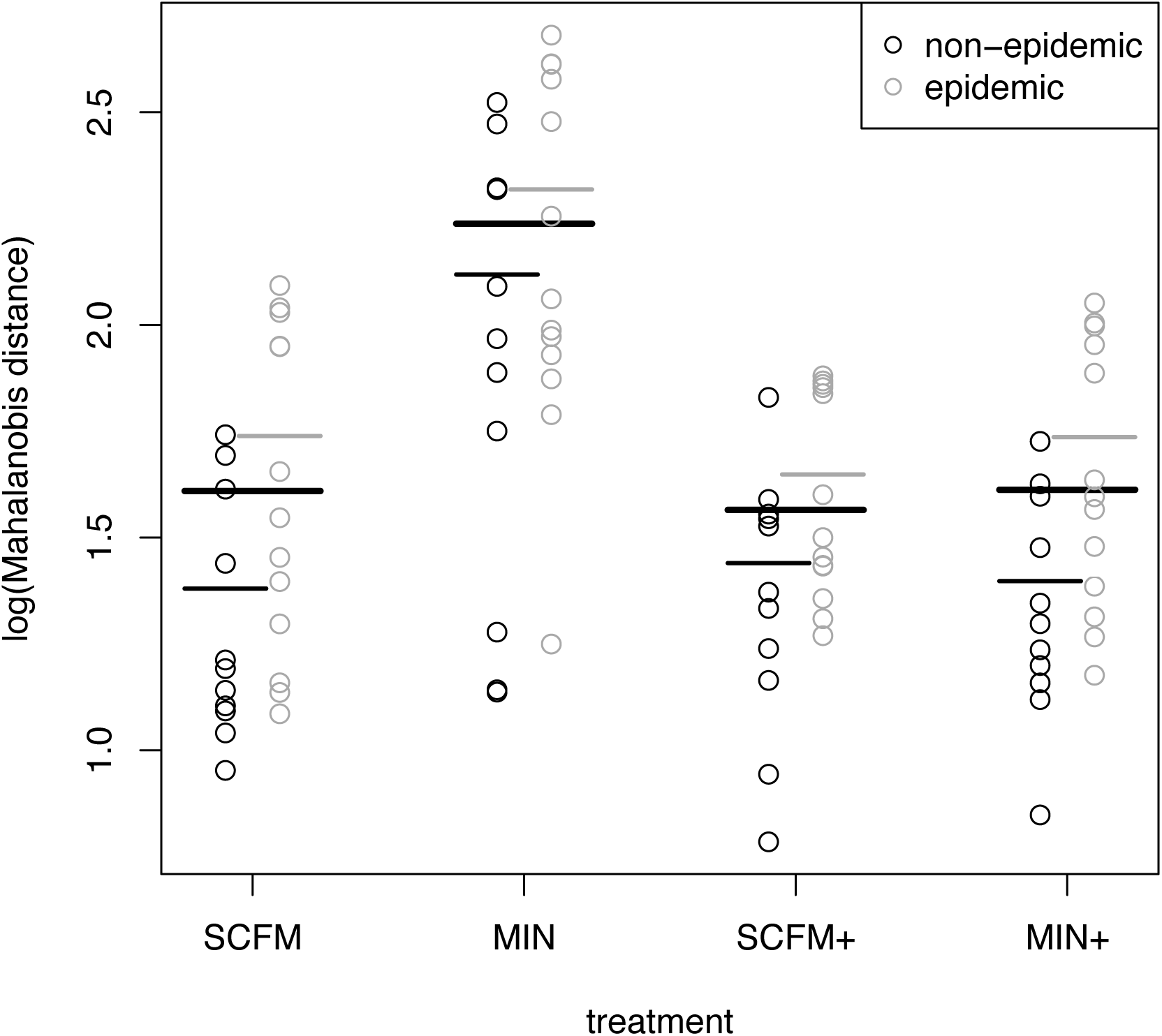
Comparison of clinical isolates to lab-evolved isolated. Each point represents the multi-variate (Mahalanobis) distance between a clinical strain and the distribution of all individual isolates from replicate populations in each treatment. Black circles are non-epidemic clinical isolates and grey circles are epidemic isolates. Solid lines represent treatment means. Mahalanobis distances were calculated using all 10 traits, *z*-transformed as before.

### Variation in colony morphology is also linked to nutritional complexity

Colony morphologies are often used as indicators for progression to chronic infection so we assayed colonies from each evolved population for 8 morphological characteristics (pigmentation, opacity, iridescence, surface texture, margin, halo, autolysis, and small colony variant). We identified a total of 18 distinct morphotypes across all 120 populations and an average of ~ 2 morphotypes per population (range from 1 to 4; SI Appendix, Fig S4). Notably, and consistent with the results presented above, populations evolved in SCFM contained more distinct colony morphs on average (ANOVA, F_1,116_ = 7.36, p = 0.008) than any other treatment, including those with mucin (ANOVA, F_1,116_ = 0.64, p = 427). Moreover, there was little correspondence between colony morphology and the changes in the suite of patho-adaptive traits we measured, as revealed by inspection of Spearman rank correlations for all pairs of traits (SI Appendix, Fig S5). These results lend further support to the growing consensus that colony morphotypes are unreliable markers of patho-adaptation and the onset of chronic infection.

### Clinical Significance and Implications

Our results provide direct experimental evidence that phenotypic diversification of *P. aeruginosa* in the airways of CF patients can be driven by divergent selection imposed by the ecological opportunity associated with the nutritionally complex lung environment. Divergence among subpopulations can be further exaggerated by reduced dispersal resulting from the thick mucus layer associated with the CFTR defect and colonization of different airway compartments. Taken together, these results suggest that the combination of nutritional complexity and reduced dispersal alone is sufficient to drive rapid and repeatable diversification, independently of other sources of selection associated with patho-adaptation such as immune evasion, resource competition from other microbial species, or redox stress.

This interpretation must be qualified, of course, by the fact that our experiments are done *in vitro* under conditions that are a far cry from the more complex and dynamic environment of the CF lung itself. The advantage of our approach is that it affords us the opportunity to construct focused, highly replicated tests of the role of nutrient complexity and mucin in driving diversification on scales that would not otherwise be possible. The disadvantage, of course, is that our environments can only represent a crude simulacrum of the conditions actually experienced over the course of an infection. That said, it is worth noting that many of the patho-adaptive phenotypes thought to be hallmarks of chronic infection, especially loss of motility and biofilm formation, were also observed in our experiment. These changes are likely due to loss-of-function mutations, a result commonly observed in the initial stages of adaptation to novel environments in microbial selection experiments (Kassen 2014). While the detailed genetic changes underlying these phenotypes will have to await whole genome sequencing (currently underway), the striking phenotypic parallelism between adaptation *in vitro* and *in vivo* suggests that the initial stages of colonization of the CF lung can be understood as a specific instance of the more general phenomenon of adaptation to a novel environment.

There are two important implications of our results for clinical practice. First, our results provide direct evidence that CF lung-like conditions promote rapid phenotypic and colony morph diversification, a result that is in line with both longitudinal (Markussen et al 2014, Clark et al 2015) and cross-sectional studies (Workentine et al 2013, Jorth et al 2015) at both the level of both phenotype and genotype. Such rapid diversification, together with the observation that a number of hallmark patho-adaptive phenotypes evolve rapidly *in vitro,* imply that colony morphology or other phenotypic biomarkers are not reliable diagnostic traits of the transition to chronic infection. Second, rapid and extensive diversification both within and among independently evolved populations suggests that the transition to chronic infection can occur by many different phenotypic and genetic routes. The presence of genetically distinct subpopulations within the lung or among different patients complicates treatment because it means that no single therapy targeting *P. aeruginosa* is likely to be effective at clearing or managing infection for all patients. Rather, a more tailored, patient-specific approach that focuses therapy on the phenotypic and genomic properties of the strain infecting a given host may prove more useful.

The long-term fate of diversity, and its consequences for patient health, remain unclear. Longitudinal studies suggest that *P. aeruginosa* diversification can occur rapidly following colonization and can persist for decades within a single host (Markussen et al 2014), likely due to specialization of sub-populations to different conditions of growth in distinct lung compartments. The comparatively short duration of our experiment does not allow us to distinguish whether the diversity we observed was transient, being the product of high mutation supply rates generating competition among independently-arising genotypes, or stable, being supported by negative frequency dependent selection linked to resource specialization. Nevertheless, our results do lend support to the idea that the nutritional conditions of the CF lung provide a substrate that spurs strain diversification and supports higher levels of genetic variation than would be otherwise available in a more homogenous environment. It may be that this diversity provides the raw material for colonization of different airway compartments and leading to compartment-based specialization. Under this view, the transition to chronic infection is intimately tied to, and is in fact the result of, diversification. This hypothesis warrants further attention.

## Materials and methods

### Bacterial strains and growth conditions

We used strain *Pseudomonas aeruginosa* Pa14, originally isolated from a human burn wound, as the founder strain for this study. Cultures were grown overnight in Luria Bertani broth (LB) and plated on LB agar to isolate individual colonies. Replicate populations were founded from distinct, single colony isolates from these plates and grown at 37 degrees C in unshaken 24-well plates.

### Selection experiment

We propagated 30 replicate populations in each of four selection environments (120 populations in total) by daily serial transfer for approximately 220 generations. Every 24 hours, 15 uL of overnight culture was transferred into 1.5 mL of fresh liquid media. We conducted a factorial experiment involving two levels of nutrient complexity (MIN: M9 minimal salts medium supplemented with 0.7% glucose; SCFM: synthetic cystic fibrosis medium, a defined medium resembling the nutritional conditions of the CF lung; see Palmer et al 2007) crossed with two levels of viscosity (supplemented or not with 50 mg/mL of porcine mucin following Wong et al 2012).

### Morphological and phenotypic diversity of endpoint populations

Aliquots from frozen archives were first revived in their respective selection environment for 24 hours at 37 degrees C and then plated on LB plates to assess colony morphology. A single observer visually identified the number of distinct colony morphs from at least 100 colonies from each population based on eight characteristics (pigmentation, opacity, iridescence, surface texture, margin, halo, autolysis, and small colony variant). Additional phenotypes, described in detail below, were measured for each of 12 random colonies from 12 random populations of each treatment for a total of 576 isolates. All phenotype assays were performed in triplicate.

### Motility

Swimming and twitching motility were measured as described in Clark et al (2015). Briefly, swim motility was assayed by stab-inoculating a sample into 0.3% LB agar plates, incubated for 24h at 37 degrees C, and their zone of growth measured. For twitch motility, colonies were stab inoculated into 1.5% LB agar and incubated for 72h (37 degrees C for 48h then room temperature for 24h). The agar was then removed from the petri dish, and the zone of growth was measured by staining the dish with 1% crystal violet.

### Pyocyanin and pyoverdine production

Cultures were first grown in LB for 24 hours and then pigment production quantified by measuring the optical density (OD) of the supernatant at 405 nm and 695 nm for pyoverdine and pyocyanin, respectively, and standardized by cell density estimated from OD at 600 nm before centrifugation and extraction of the supernatant.

### Biofilm formation

Biofilm formation was assayed as described by O’Toole (2011). Cultures were diluted 1:100 in fresh LB, seeded into 4 replicate wells of a 96-well non-tissue culture treated microtitre plate and incubated for 24 h at 37 degrees C under static conditions. Plates were then rinsed and stained with 125 uL of a 0.1% solution of crystal violet in water. After incubating at room temperature for 15 mins, the plates were rinsed several times with water and allowed to dry for 48 hours. The stain was then dissolved in 125 uL of 30% acetic acid in water and incubated at room temperature for 15 mins. Biofilm formation was quantified by measuring the optical density of this final solution at 550nm.

### Antibiotic resistance

We used a two-fold dilution series for each of four antibiotics (ciprofloxacin, ceftazidime, tobramycin, and colistin) in LB media to estimate minimum inhibitory concentration (MIC; the lowest concentration of drug that prevents growth) for each evolved isolate and the founding strain. Growth was assessed at 24h and 48h by measuring optical density at 600 nm and the MIC determined as the drug concentration where OD was less than 0.100. In cases where growth was observed at all concentrations of antibiotic, the MIC was reported, conservatively, as the next highest concentration in the dilution series.

### Statistical analysis

All statistical analyses were performed in R, version 3.3.0 (R Core Team, 2017). All phenotypic traits were treated as continuous variables, with antibiotic resistance traits analyzed on a logarithmic scale. To determine the effect of media, mucin, and the interaction on each phenotype individually, we used the R package nlme to construct linear mixed effects models with population modeled as a random effect and media and mucin as fixed effects. We performed *post hoc* pairwise *t*-tests, Holm-adjusted for multiple comparisons, to evaluate the statistical significance among treatments. Trait values of evolved populations were compared to those of the unevolved founder using one-sample *t*-tests within each treatment and corrected for multiple comparisons (40 in total) using a Bonferroni corrected *P*-value of α = 0.00125.

Diversity within a population was determined by calculating the Euclidean distance on standardized z-scores for each trait between all possible pairs of isolates. Diversity between populations was determined by calculating the Euclidean distance between the means of all populations within a treatment group for all possible pairs of populations. A randomization test was performed to determine significance by resampling treatment labels and calculating a null distribution of F-statistics. To determine the distance between the clinical strains and the evolved isolates, we calculated the Mahalanobis distance between each clinical strain and the distribution of treatment group. Correlations between phenotypes were determined by calculating Spearman correlation coefficients and visualized using the package ‘gplots’ in R.

## Supplementary Figures and Tables

**Supplementary Table 1.** Treatment effects on individual phenotypic traits. Values are from linear mixed effects models with media (MIN or SCFM) and mucin (present or absence) as fixed effects and population as a random effect. P-values < 0.05 are in bold.

| Trait | media |  | mucin |  | media * mucin |  |
| --- | --- | --- | --- | --- | --- | --- |
|  | F<sub>(1,44)</sub> ratio | P-value | F<sub>(1,44)</sub> ratio | P-value | F<sub>(1,44)</sub> ratio | P-value |
| Growth rate | 0.56 | 0.457 | 0.90 | 0.347 | 6.25 | **0.016** |
| Pyoverdine | 0.05 | 0.832 | 11.56 | **0.001** | 0.03 | 0.875 |
| Pyocyanin | 0.80 | 0.375 | 3.47 | 0.069 | 2.88 | 0.097 |
| Biofilm | 13.75 | **< 0.001** | 25.93 | **< 0.001** | 25.29 | **< 0.001** |
| Swim mot. | 0.17 | 0.679 | 66.18 | **< 0.001** | 1.73 | 0.196 |
| Twitch mot. | 31.16 | **< 0.001** | 0.85 | 0.362 | 0.01 | 0.917 |
| MIC cip | 31.07 | **< 0.001** | 4.19 | **0.047** | 25.29 | **< 0.001** |
| MIC cef | 0.36 | 0.553 | 7.36 | **0.010** | 1.54 | 0.222 |
| MIC col | 1.10 | 0.300 | 0.10 | 0.756 | 0.58 | 0.452 |
| MIC tob | 1.73 | 0.195 | 0.40 | 0.531 | 0.27 | 0.608 |

**Supplementary Figure 1.**
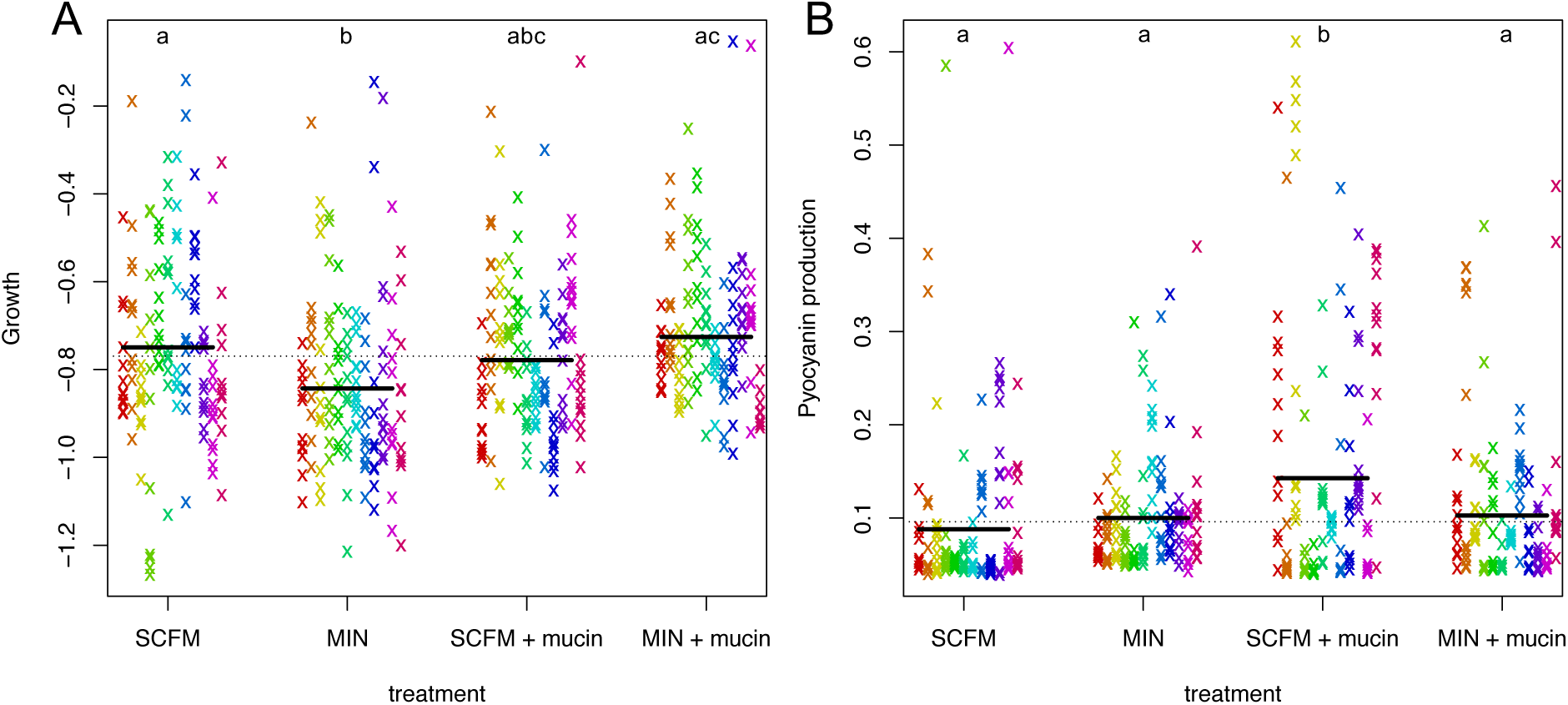
Phenotypic adaptation in evolved isolates for growth (A) and pyocyanin production (B). Within each treatment, each column/colour represents a population. Solid lines represent treatment means and dashed lines represent the value of that phenotype in the ancestral strain (Pa14).

**Supplementary Figure 2.**
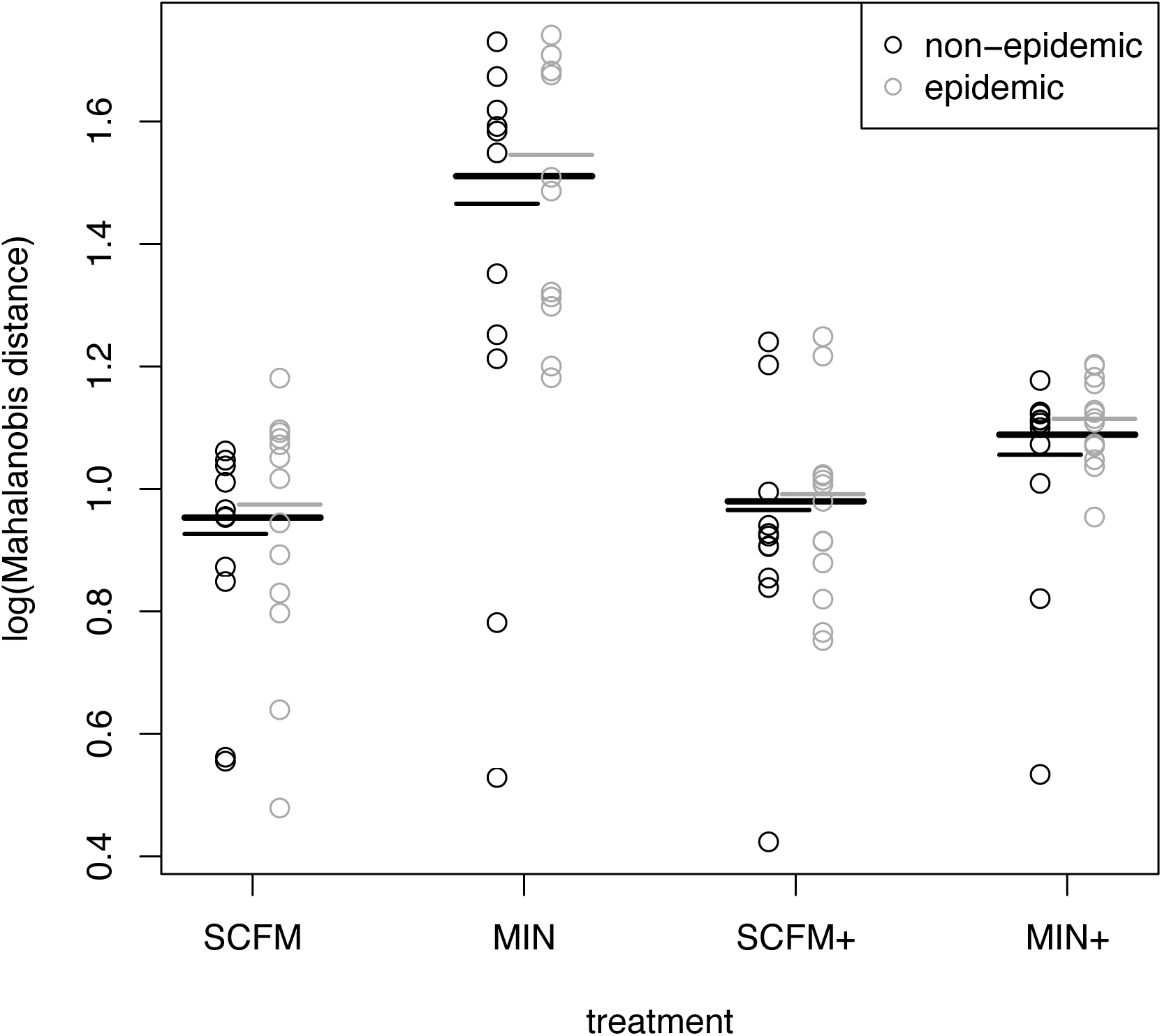
Comparison of clinical isolates to lab-evolved isolated. Each point represents the multi-variate (Mahalanobis) distance between a clinical strain and the distribution of all individual isolates within that treatment. Black circles are non-epidemic clinical isolates and grey circles are epidemic isolates. Solid lines represent treatment means. Here, Mahalanobis distance was calculated using all trait measurements excluding antibiotic resistance traits.

**Supplementary Figure 3.**
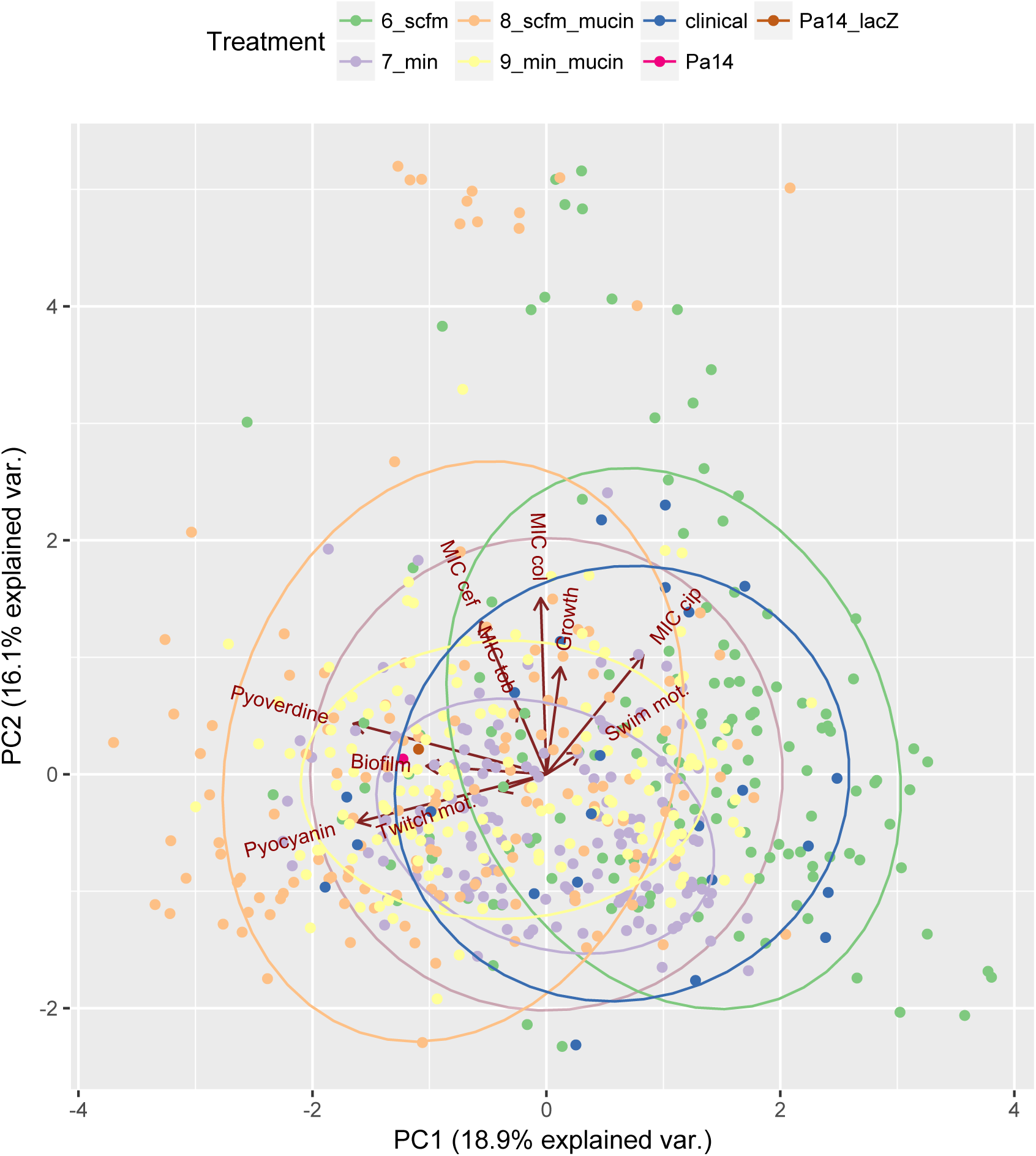
Principal component analysis of all evolved isolates and clinical isolates. Each point represents an individual and each treatment is grouped. Ancestral strains, Pa14 and Pa14_lacz, are single pink and brown points, respectively.

**Supplementary Figure 4.**
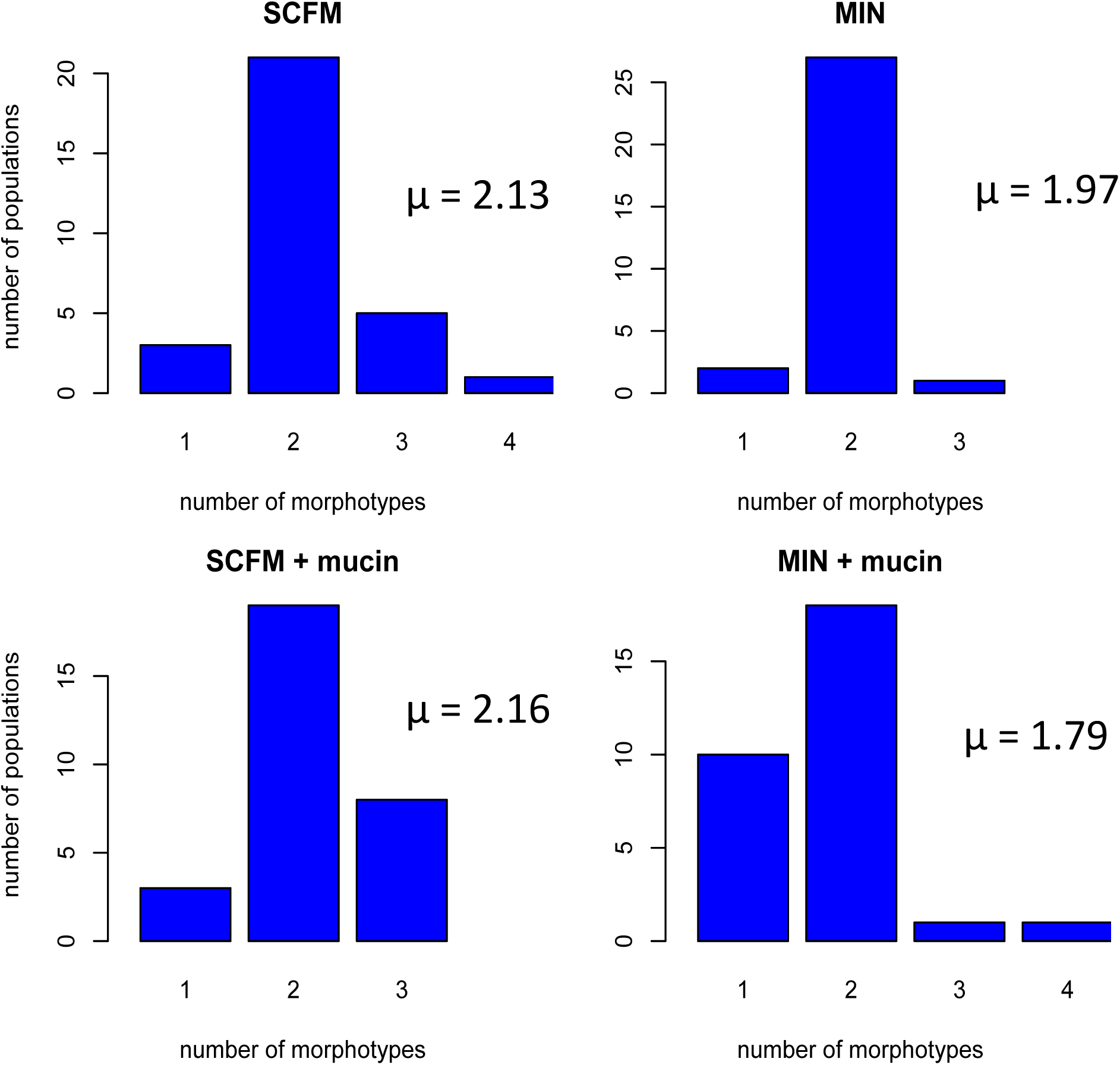
Number of distinct morphologies present in all evolved populations. Colonies were categorized using 8 characteristics: pigmentation, opacity, iridescence, surface texture, margin, halo, autolysis, and small colony variant.

**Supplementary Figure 5.**
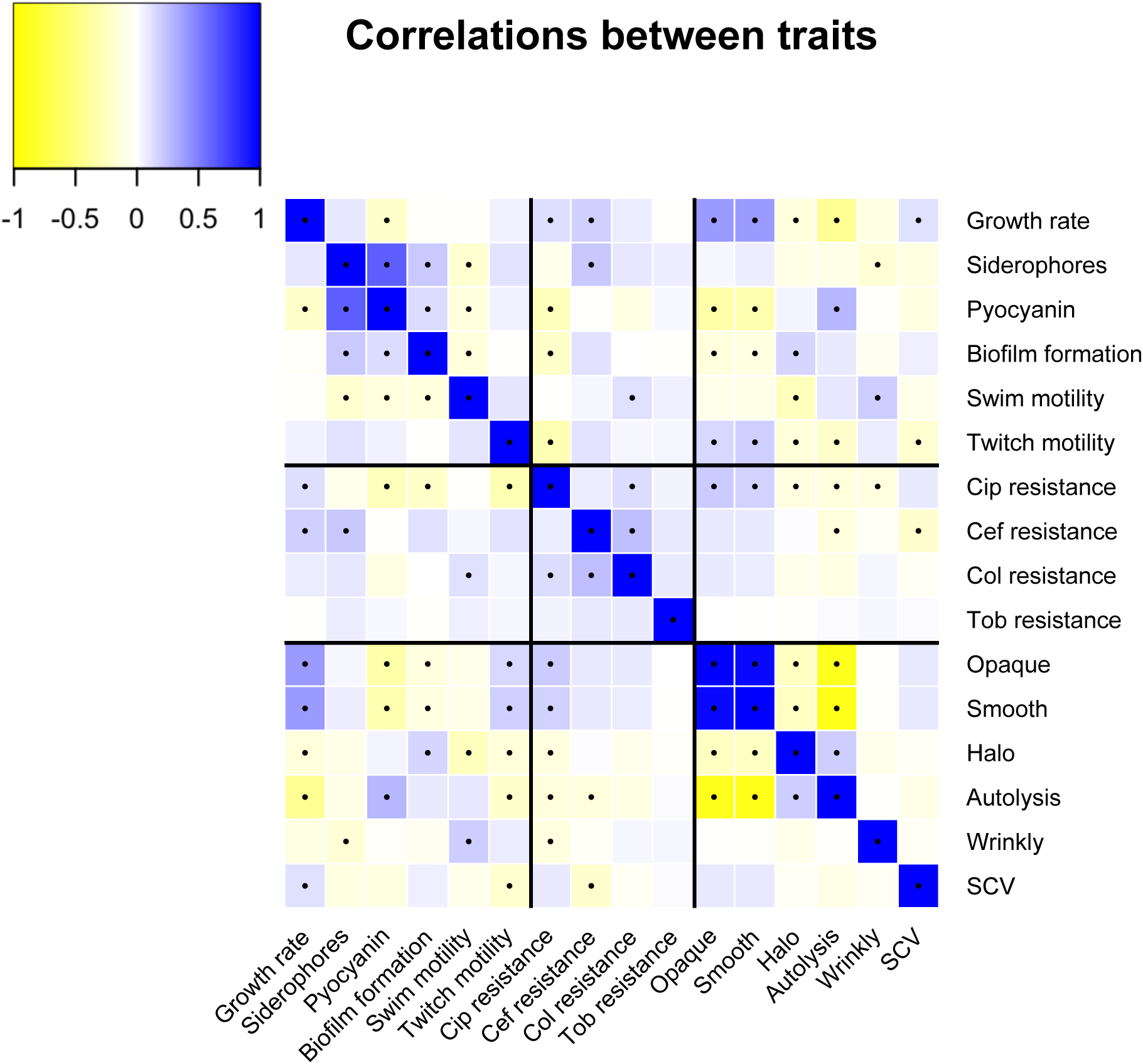
Spearman rank correlations between all traits for all evolved isolates. Black lines separate groups of traits, divided into phenotypic traits, antibiotic-resistance, and colony morphology traits. Yellow indicates strong positive correlation between any pair of traits and blue indicates strong negative correlation. Black dots within boxes represent a statistically significant correlation (p < 0.05).

